## Supplementary Figs. 1-10 for "Inhibiting Stromal Class I HDACs Curbs Pancreatic Cancer Progression"

Liang *et al. Nature Communications.*

### SUPPLEMENTARY INFORMATION

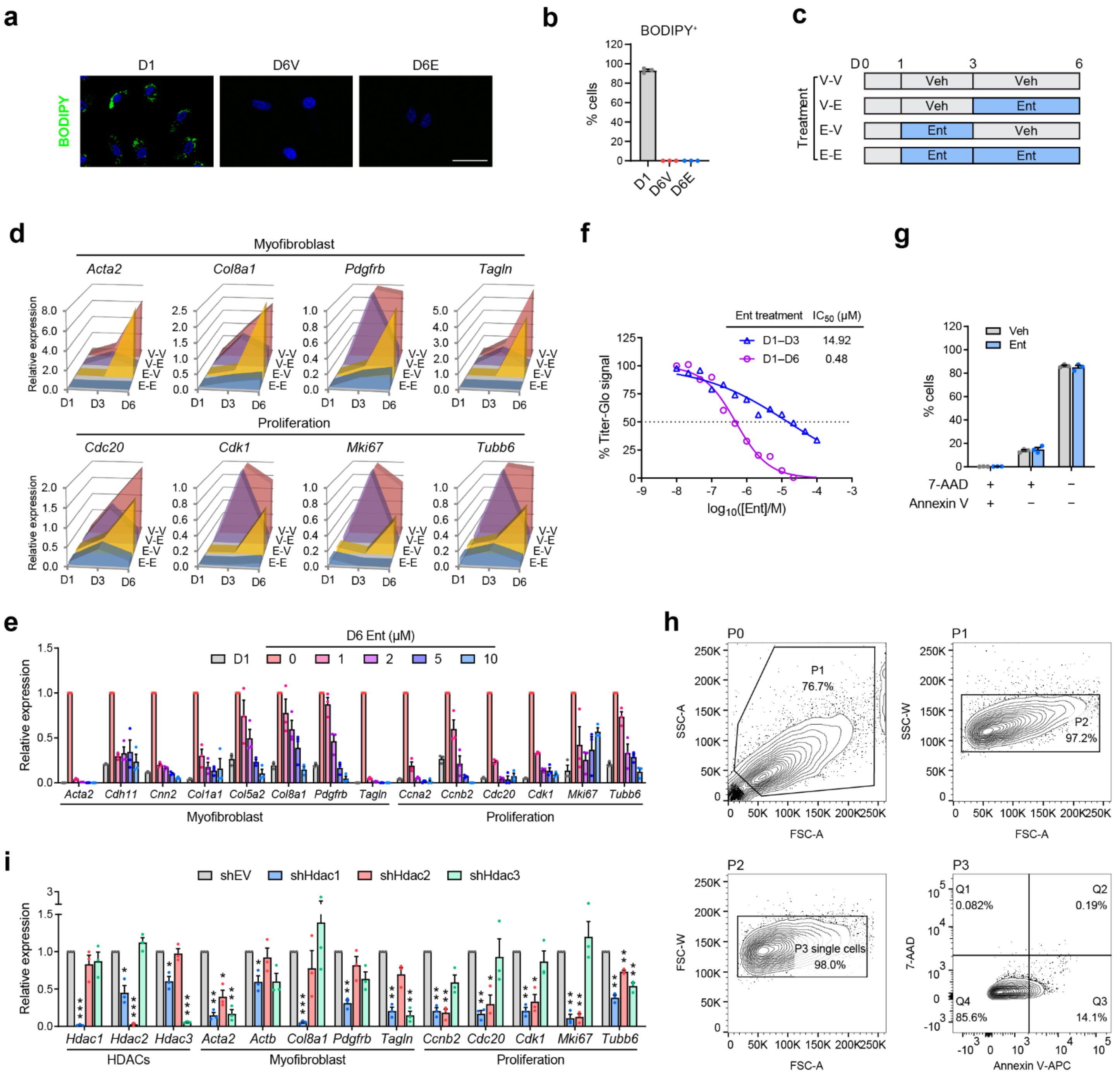


**Supplementary Fig. 1 | Effects of HDACi on PSC activation.**

**a, b** Representative images (**a**) and quantifications (**b**) of lipid content staining by BODIPY in PSC samples. $n$ = 3 independent samples. Scale bar, 50 μm. **c** Periodic treatment scheme of Ent during PSC activation. **d** Representative RT-qPCR data showing the expression of representative markers for myofibroblast identity and proliferation in PSC samples under periodic treatments (**c**), compared to Veh treatment at D3. 3 independent experiments were performed with similar results. **e** RT-qPCR data showing the expression of myofibroblast and proliferation genes in PSCs at D1 and D6 with or without 5-day Ent treatments (1–10 μM). $n$ = 3 independent samples. **f** Dosage response curves with $IC_{50}$ from Titer-Glo assays in PSCs treated with Ent from D1 to D3 or D6. $n$ = 5 cell sample replicates. Mean values are presented. **g** Results from flow cytometry with staining of apoptotic markers Annexin V and 7-AAD in PSCs under Veh or Ent treatment (5 μM) for 5 d. $n = 3$ cell sample replicates. **h** Gating scheme for Annexin V and 7-AAD analysis from bulk population (P0) to subpopulations with differential Annexin V and 7-AAD staining (Q1–4). **i** RT-qPCR results showing the expression of HDACs and selected PSC activation markers upon depletions of individual HDACs by shRNAs, compared to shEV. $n = 3$ independent samples. Data in bar graphs are presented as mean values ± SEM. *, $p < 0.05$; **, $p < 0.01$; ***, $p < 0.001$. Two-sided t-test.

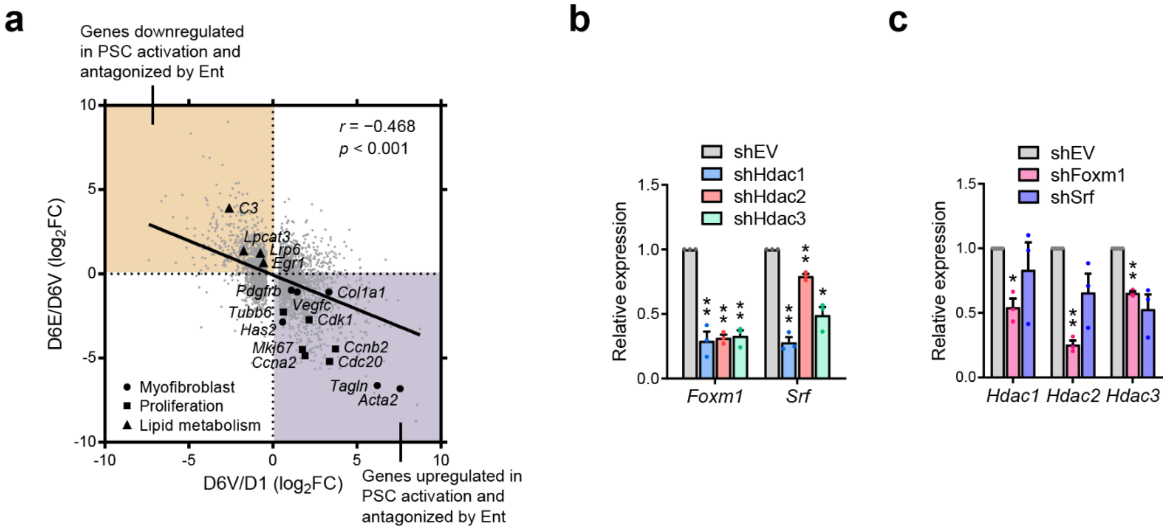


**Supplementary Fig. 2 | Transcriptional control of PSC activation by HDACs and TFs.**

**a** Scatter plot from RNA-seq data showing the reverse correlation between the expression changes in PSC activation and those under Ent treatment at genes significantly upregulated or downregulated in PSC activation, including representative genes functioning in myofibroblast (circle), proliferation (square) and lipid metabolism (triangle). **b, c** RT-qPCR data showing the expression of TFs in activated PSCs with shRNAs depleting individual HDACs (**b**), and the expression of HDACs in those with shRNAs depleting individual TFs (**c**). RNA-seq and RT-qPCR, $n = 3$ independent samples. Data in bar graphs are presented as mean values ± SEM. *, $p < 0.05$; **, $p < 0.01$. Two-sided t-test.

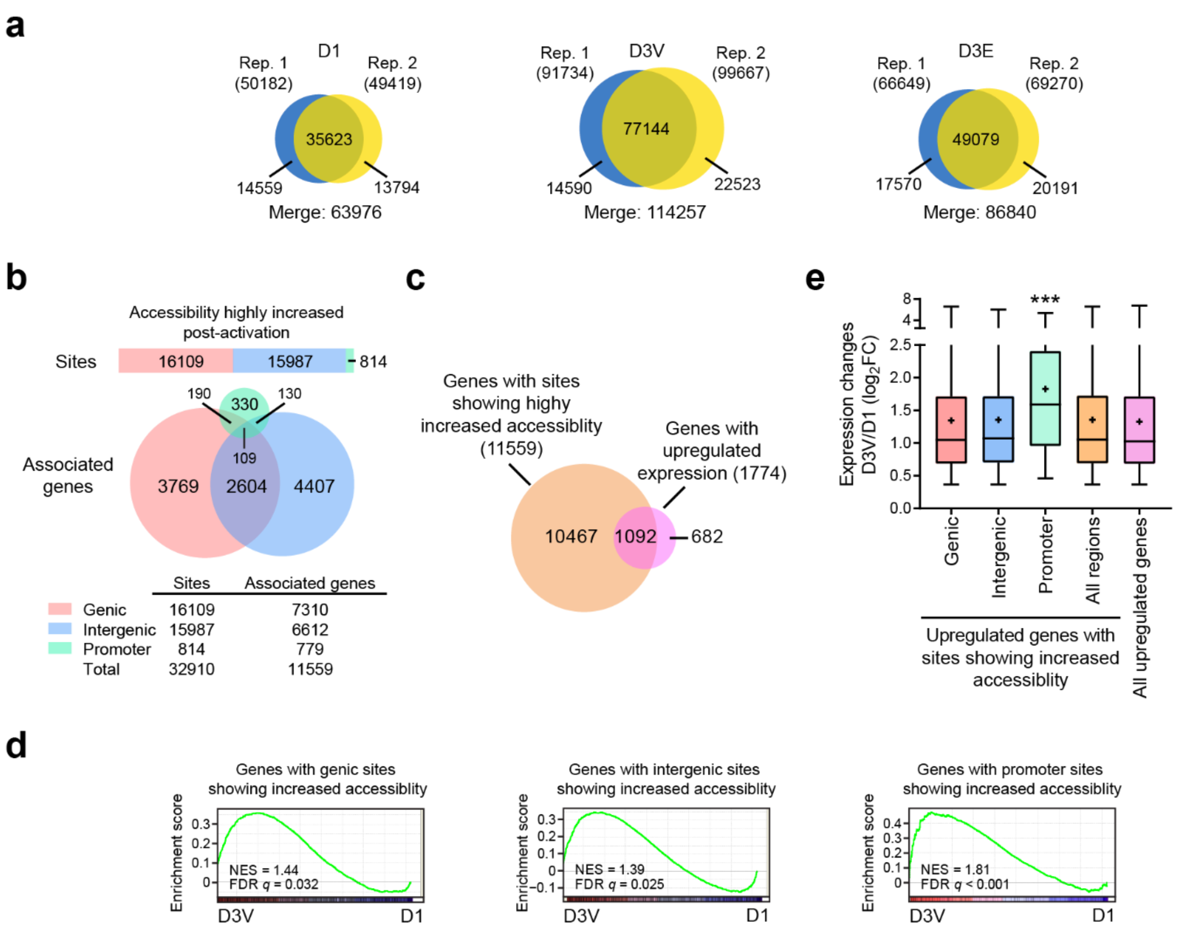


**Supplementary Fig. 3 | Genomic sites with highly increased accessibility are linked to transcriptional upregulation in activated PSCs.**

**a** Venn diagrams showing the distributions of accessible genomic sites in each ATAC-seq replicate (Rep.) of the PSC samples. **b** Numbers of genomic sites with highly increased accessibility post-activation (D3V vs D1; FC > 4, FDR $q < 0.05$) and their associated genes. **c** Venn diagram showing the distribution of genes associated with genomic sites showing highly increased accessibility and genes significantly upregulated post-activation. **d** GSEA plots showing the enrichment of genes upregulated in PSC activation in the gene sets linked to genic, intergenic or promoter sites showing highly increased accessibility post-activation. **e** Box plots showing the expression inductions of gene sets upregulated post-activation and linked to genomic sites showing increased accessibility at different annotated regions. Center lines, medians; box limits, first and third quartiles; whiskers, minima (first quantiles − 1.5 × IQR) and maxima (third quantiles + 1.5 × IQR); +, mean. IQR, interquartile range. ***, $p < 0.001$, promoter compared to any other groups. One-way ANOVA.

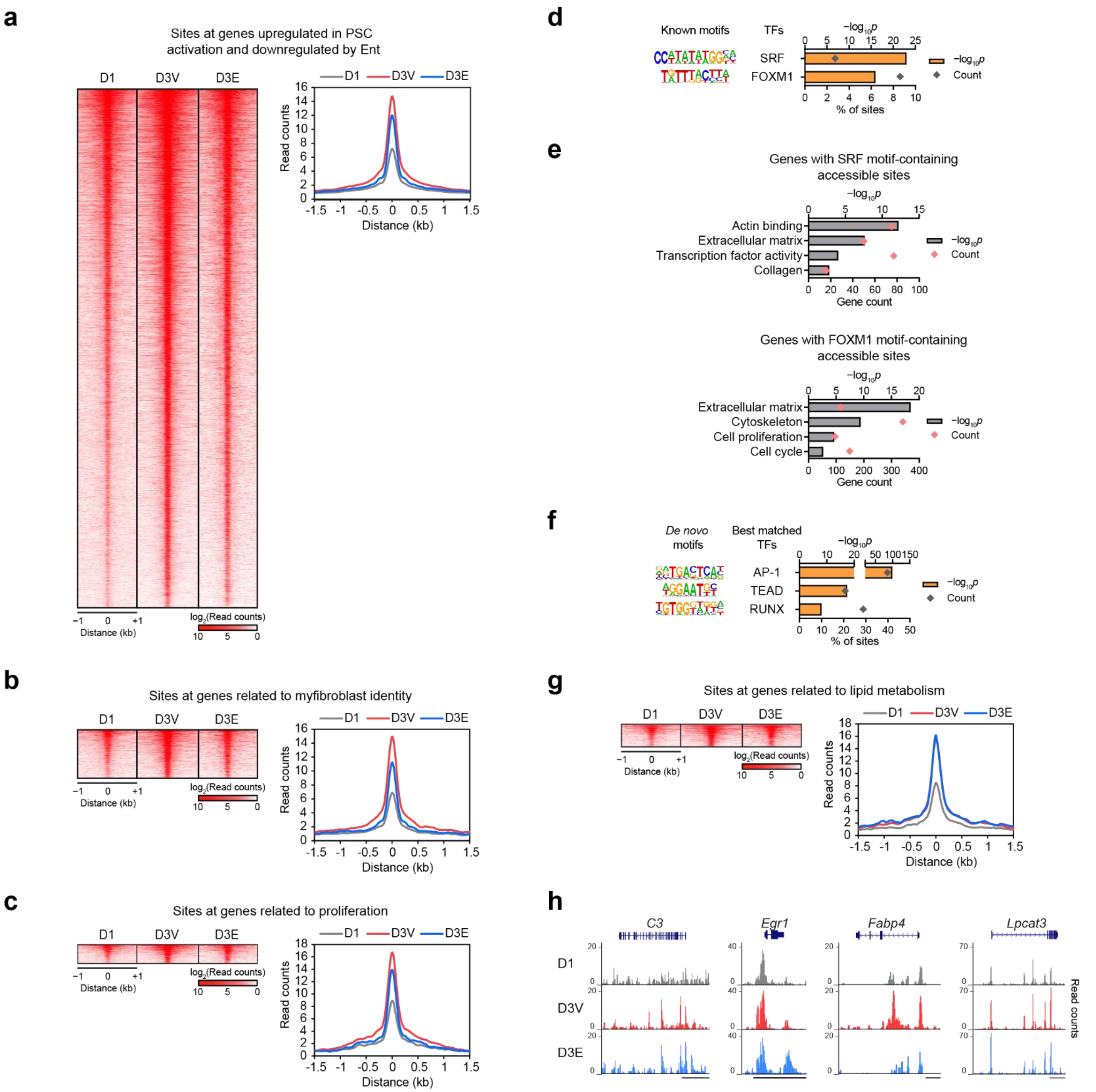


**Supplementary Fig. 4 | Genomic accessibility of functional gene sets in PSC activation.**

**a–c** Histograms and heatmaps showing the accessibility of genomic sites associated with selected gene sets, including genes upregulated in PSC activation and downregulated by Ent (**a**), as well as subsets of genes related to myofibroblast identity (**b**) and proliferation (**c**). **d** Selected enriched TF motifs at genomic sites with significantly increased accessibility post-activation. **e** Representative ontology terms for gene sets associated with genomic sites containing SRF or FOXM1 motif and showing increased accessibility post-activation. **f** Top *de novo* motifs and their best matched TFs identified from genomic sites with significantly increased accessibility post-activation. **g** Histograms and heatmaps showing the accessibility of genomic sites at or near lipid metabolism-related genes that are downregulated by PSC activation and upregulated by Ent. **h** Genome browser tracks for selected lipid metabolism-related genes. Scale bar, 1 (*Fabp4*) or 10 kb (others). Motif (**d, f**) and GO analyses (**e**), one-sided Fisher's exact test. Genes included in the analyses in **b**, **c**, **g** are listed in Supplementary Data 2.

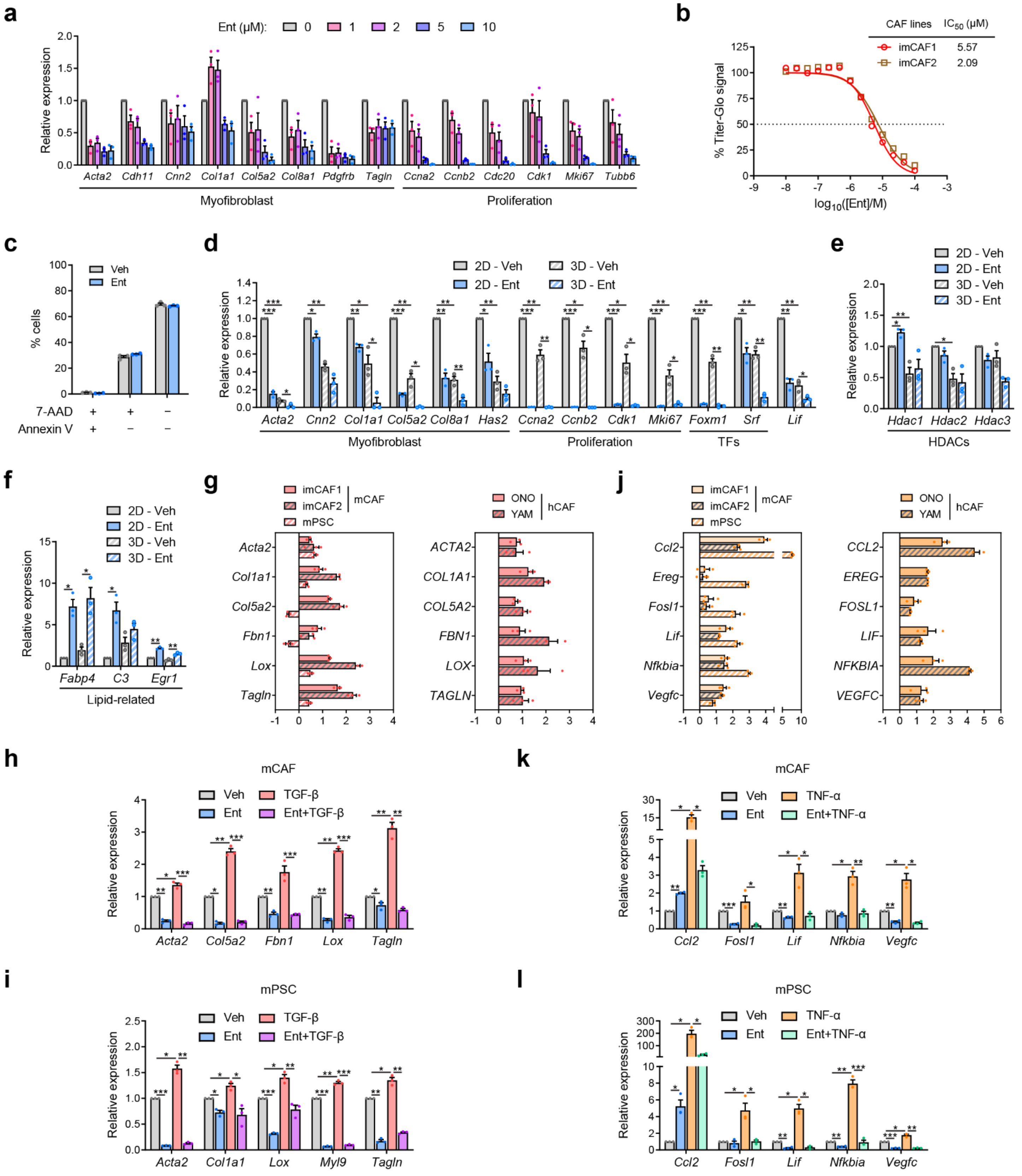


**Supplementary Fig. 5 | CAF responses to Ent, 3D culture and cytokine stimulations.**

**a** Gene expression detected by RT-qPCR in mCAFs (imCAF1) under Ent treatment (2 d) with selected concentrations compared to no Ent treatment. *n* = 3 independent samples. **b** Dosage response curves with $IC_{50}$ from Titer-Glo assays of Ent-treated mCAFs. *n* = 5 cell sample replicates. Mean values are presented. **c** Data from flow cytometry with Annexin V and 7-AAD staining in imCAF1 under Veh or Ent treatment (10 μM, 2 d). *n* = 3 cell sample replicates. **d–f** RT-qPCR results showing the expression of representative Ent-downregulated myofibroblast, proliferation and TF genes (**d**), HDAC genes (**e**) and representative Ent-upregulated lipid-related genes (**f**) in mouse CAFs (imCAF1) in 2D or 3D culture after Veh or Ent treatment (10 μM, 2 d). *n* = 3 independent samples. **g** Bar graphs showing the expression FC measured by RT-qPCR at representative TGF-β-induced genes in mCAF (imCAF1, imCAF2), mPSC and hCAF (ONO, YAM) cultures upon TGF-β treatment (1 ng/ml, 2 d). *n* = 3 independent samples. **h, i** RT-qPCR data showing the effects of Ent on the expression of TGF-β-induced genes in mCAF (imCAF1, **h**) and mPSC (**i**) cultures, compared to Veh. *n* = 3 independent samples. **j** Bar graphs showing the expression FC measured by RT-qPCR at representative TNF-α-induced genes in mCAF, mPSC and hCAF cultures upon TNF-α treatment (10 ng/ml, 8 h). *n* = 3 independent samples. **k, l** RT-qPCR data showing the effects of Ent on the expression of TNF-α-induced genes in mCAF (imCAF1, **k**) and mPSC (**l**) cultures, compared to Veh. *n* = 3 independent samples. Data in bar graphs are presented as mean values ± SEM. *, $p < 0.05$; **, $p < 0.01$; ***, $p < 0.001$. Two-sided t-test.

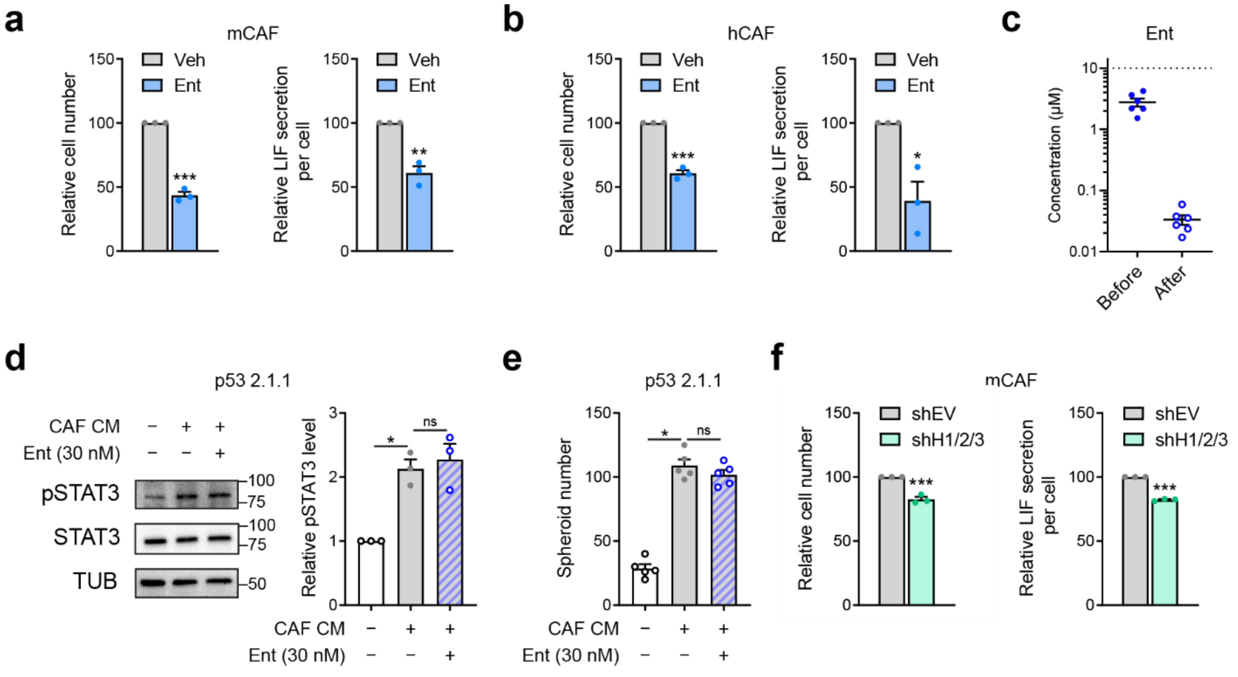


**Supplementary Fig. 6 | Analyses of relative LIF secretion per cells and the effects of residual Ent in CM.** **a, b** Relative cell numbers in mCAF (imCAF1, **a**) and hCAF (YAM, **b**) cultures at CM harvest and estimated relative LIF secretion per cells after Veh or Ent (10 µM, 2 d) treatment. *n* = 3 independent samples. **c** Ent concentrations measured by mass spectrometry in CM before and after small molecule depletion, compared to initially applied concentration (dotted line). *n* = 6 independent samples. **d** Representative images and quantifications of pSTAT3/STAT3 ratio from Western blotting with p53 2.1.1 cells treated with CM in the presence or absence of the residual level of Ent (30 nM). *n* = 3 independent samples. Uncropped blot images in Supplementary Information. **e** Spheroid numbers from p53 2.1.1 cells under the treatment of CM with or without Ent (30 nM). *n* = 5 cell sample replicates. **f** Relative cell numbers in mCAF (imCAF1) cultures with shEV or shH1/2/3 at CM harvest, and estimated relative LIF secretion per cells. *n* = 3 independent samples. Data are presented as mean values ± SEM. *, $p < 0.05$; **, $p < 0.01$; ***, $p < 0.001$; ns, not significant. Two-sided t-test.

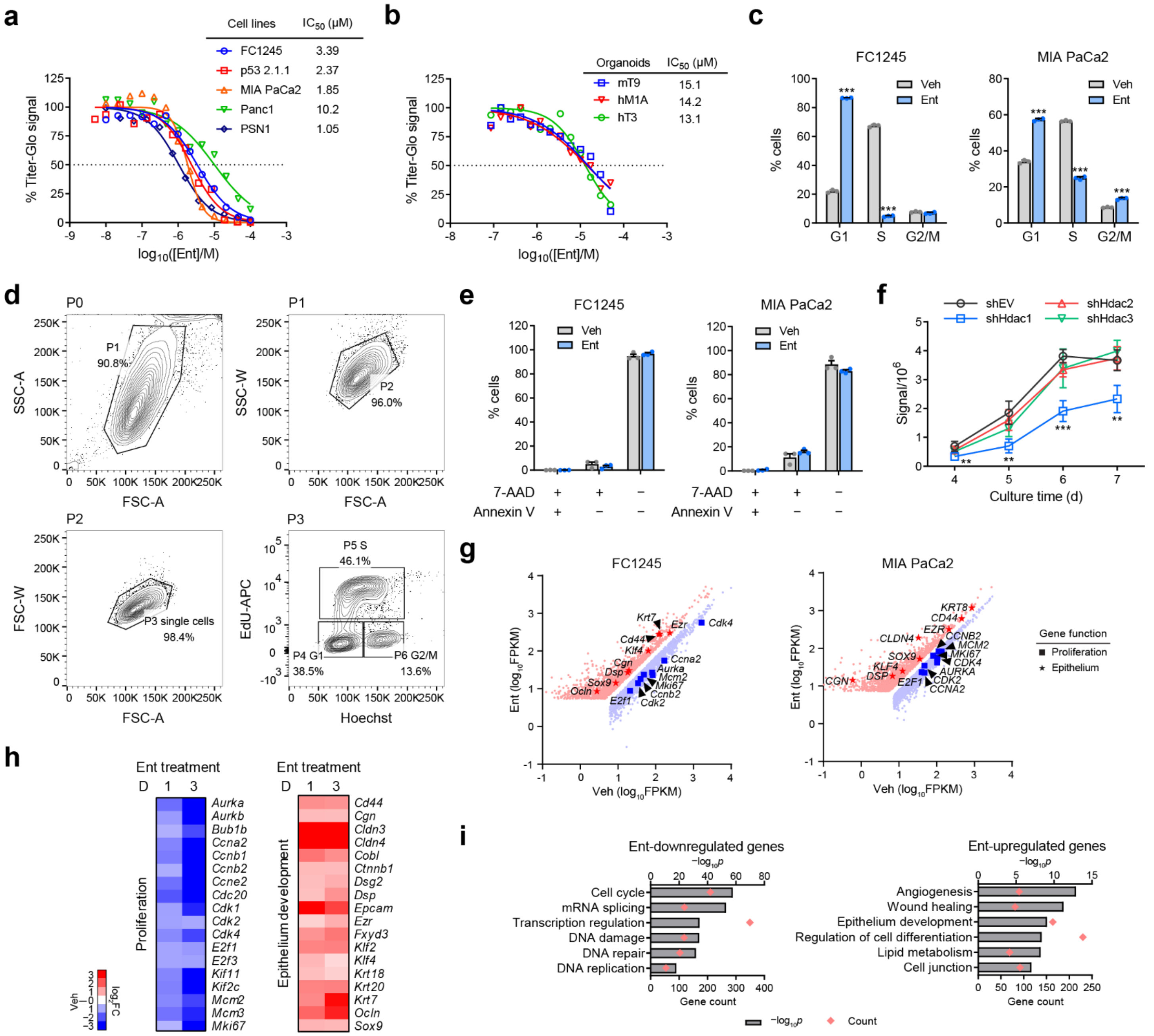


**Supplementary Fig. 7 | In vitro Effects of HDACi on PDAC cells and organoids.**

**a, b** Dose response curves with $IC_{50}$ from Titer-Glo assays with Ent-treated PDAC cells and organoids subjected to Ent treatment (**a**; 2 d for p53 2.1.1, 3 d for others) and organoids (**b**; 3 d). *n* = 5 (FC1245, MIA PaCa2, Panc1) and 3 (p53 2.1.1, PSN1, all organoids) cell sample replicates. Mean values are presented. **c** Results from EdU incorporation assay showing the cell cycle distribution of mouse (FC1245) and human PDAC cells (MIA PaCa2) under Veh or Ent treatment (5 μM, 2 d). *n* = 3 cell sample replicates. **d** Gating scheme for EdU assay from bulk population (P0) to G1 (P4), S (P5) and G2/M (P6) populations. **e** Results from flow cytometry for Annexin V and 7-AAD staining in FC1245 and MIA PaCa2 cells under Veh or Ent treatment (5 μM, 2 d). *n* = 3 cell sample replicates. **f** GFP signals detected in GFP-labelled FC1245 cells transduced with shRNAs against individual HDACs along 7-day culture, compared to shEV. *n* = 5 cell sample replicates. shEV vs shHdac1, $p = 0.007$ (D4), 0.001 (D5, D7), $p < 0.001$ (D6). Two-sided t-test. **g** Scatter plots from RNA-seq data showing genes significantly upregulated (red) and downregulated (blue) by Ent treatments (5 μM, 1

d) in FC1245 and MIA PaCa2 with selected markers for proliferation and epithelium highlighted. $n = 2$ independent samples. **h** Heatmap showing the expression change of selected Ent-downregulated genes functioning in proliferation and selected Ent-upregulated genes functioning in epithelium development in FC1245 after 1 or 3 d Ent treatment. **i** Selected ontology terms enriched in gene sets downregulated or upregulated by Ent in FC1245. GO analysis, one-sided Fisher's exact test. Data in bar and line graphs are presented as mean values ± SEM. **, $p < 0.01$, ***, $p < 0.001$. Two-sided t-test.

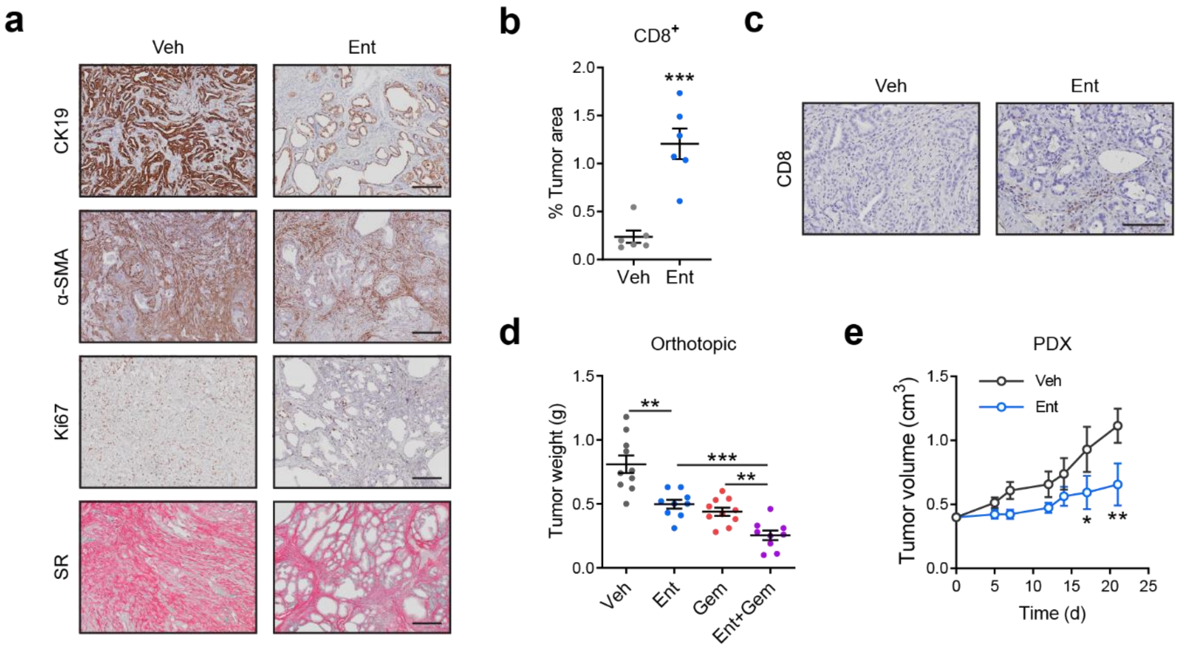


**Supplementary Fig. 8 | Effects of Ent treatment in PDAC mouse models.**

**a** Representative views of CK19, α-SMA and Ki67 staining in tumor sections from $KP^{f/f}C$ mice with Veh or Ent treatment. Scale bar, 200 μm. **b, c** Quantifications of the percentage of $CD8^+$ staining in tumor area (**b**) and representative images of $CD8^+$ T cells (**c**) from tumor sections of $KP^{f/f}C$ mice with Veh or Ent treatment. Scale bar, 100 μm. $n = 6$ tumors. **d** Tumor weights from syngeneic orthotopic transplants of mouse PDAC cells (FC1245) after 2-week treatment of Veh, Ent (15 mg/kg, daily), Gem (5 mg/kg, q3d) or the combination of Ent and Gem. $n = 10$ (Veh, Gem) and 9 (Ent, Ent+Gem) mice. **e** Tumor progression curves showing the tumor volumes of subcutaneously implanted PDX in athymic mice treated with Veh or Ent (10 mg/kg, daily). $n = 6$ mice. *, $p < 0.05$; **, $p < 0.01$; ***, $p < 0.001$.Data are presented as mean values ± SEM. Two-sided t-test.

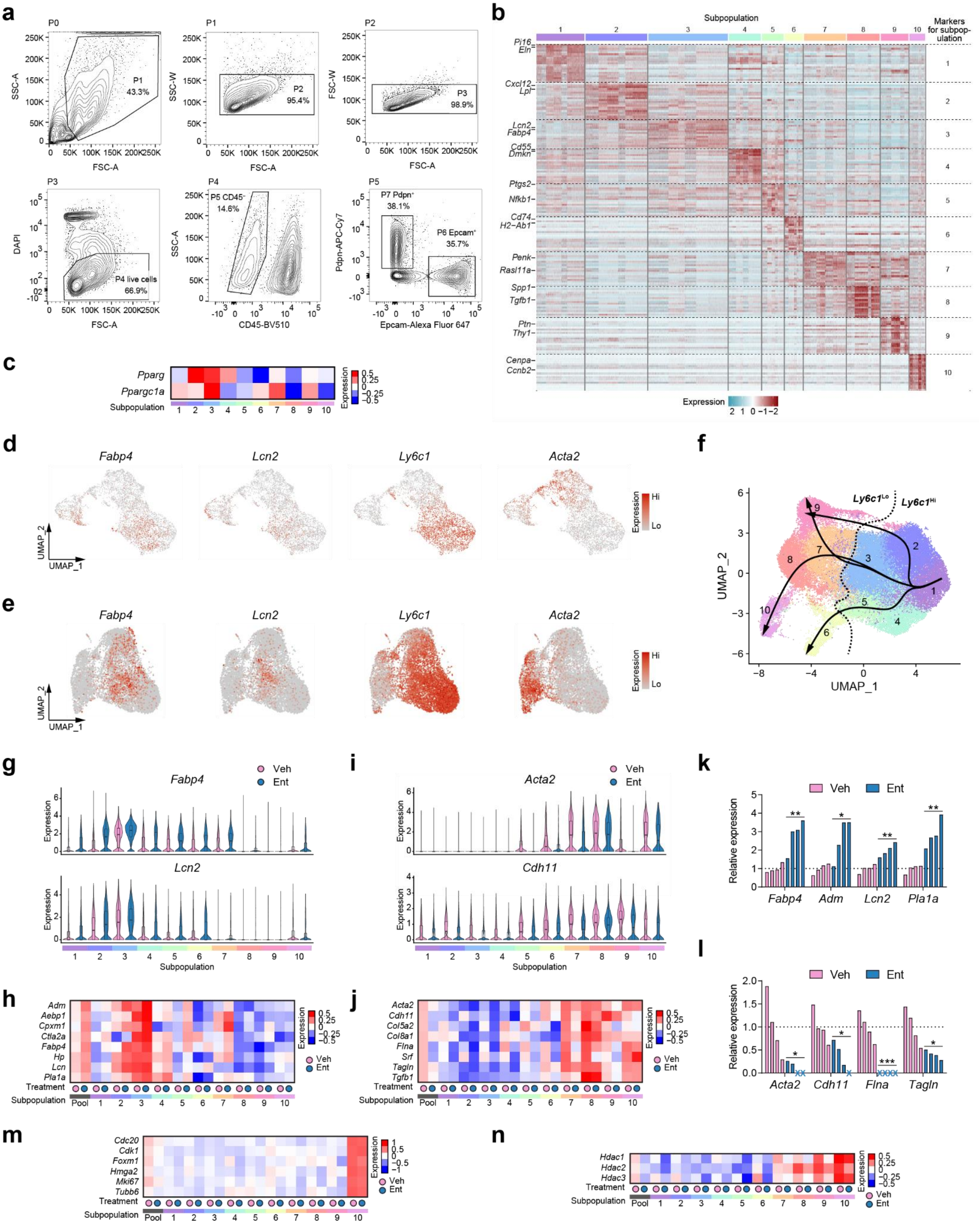
a
P0
P1 43.3%
SSC-A
FSC-A
P1
P2 95.4%
SSC-W
P2
P3 98.9%
FSC-W
P3
P4 live cells 66.9%
DAPI
P4
P5 CD45- 14.6%
CD45-BV510
P5
P7 Pdpn+ 38.1%
P6 Epcam+ 35.7%
Pdpn-APC-Cy7
Epcam-Alexa Fluor 647
b
Subpopulation
Markers for subpopulation
Pi16
Eln
Cxcl12
Lpl
Lcn2
Fabp4
Cd55
Dmkn
Ptgs2
Nfkb1
Cd74
H2-Ab1
Penk
Rasl11a
Spp1
Tgfb1
Ptn
Thy1
Cenpa
Ccnb2
Expression
c
Pparg
Ppargc1a
Subpopulation
Expression
d
Fabp4
Lcn2
Ly6c1
Acta2
UMAP_2
UMAP_1
Expression
Hi
Lo
e
Fabp4
Lcn2
Ly6c1
Acta2
f
Ly6c1Lo
Ly6c1Hi
UMAP_2
UMAP_1
g
Fabp4
Lcn2
Veh
Ent
Expression
Subpopulation
i
Acta2
Cdh11
Veh
Ent
Expression
Subpopulation
k
Veh
Ent
Relative expression
Fabp4
Adm
Lcn2
Pla1a
l
Veh
Ent
Relative expression
Acta2
Cdh11
Flna
Tagln
h
Adm
Aebp1
Cpxm1
Ctla2a
Fabp4
Hp
Lcn
Pla1a
Treatment
Subpopulation
Pool
Expression
Veh
Ent
j
Acta2
Cdh11
Col5a2
Col8a1
Flna
Srf
Tagln
Tgfb1
Treatment
Subpopulation
Pool
m
Cdc20
Cdk1
Foxm1
Hmga2
Mki67
Tubb6
Treatment
Subpopulation
Pool
n
Hdac1
Hdac2
Hdac3
Treatment
Subpopulation
Pool

**Supplementary Fig. 9 | Transcriptional effects of Ent on stromal fibroblasts at a single-cell level.**

**a** Gating scheme for isolating stromal fibroblasts (P7, Pdpn$^+$) from the bulk cell population (P0) of dissociated tumors from *KP$^{f/f}$C* mice. **b** Heatmap showing the expression of the top 25 markers identified in each fibroblast subpopulation. **c** Heatmap showing the expression of lipogenic regulators in fibroblast subpopulations. **d, e** UMAP showing the expression of *Fabp4*, *Ly6c1* and *Acta2* in stromal fibroblast populations in scRNA-seq datasets from Elyada *et al.*[1] (**d**) and Buechler *et al.*[2] (**e**). **f** Trajectories inferred by Slingshot analysis across fibroblast subpopulations. **g–j** Violin plots and heatmaps showing the expression of lipogenic (**g, h**) and myofibroblastic (**i, j**) markers in the fibroblast subpopulations from Veh and Ent samples. Boxes in violin plots (**g, i**): center lines, medians; box limits, first and third quartiles; whiskers, minima (first quantiles − 1.5 × IQR) and maxima (third quantiles + 1.5 × IQR). IQR, interquartile range. **k, l** Waterfall plots from RT-qPCR data showing the expression of lipogenic (**k**) and myofibroblastic (**l**) markers in stromal fibroblasts from the Ent samples relative to the Veh samples. $n$ = 4 mouse tumors. Dotted lines, average expression in Veh; crosses, samples with undetectable expression. *, $p < 0.05$; ** $p < 0.01$; ***, $p < 0.001$. Two-sided t-test. **m, n** Heatmaps showing the expression of *Foxm1* and other cell cycle-related genes (**m**), and HDAC genes (**n**) in fibroblast subpopulation in Veh and Ent samples.

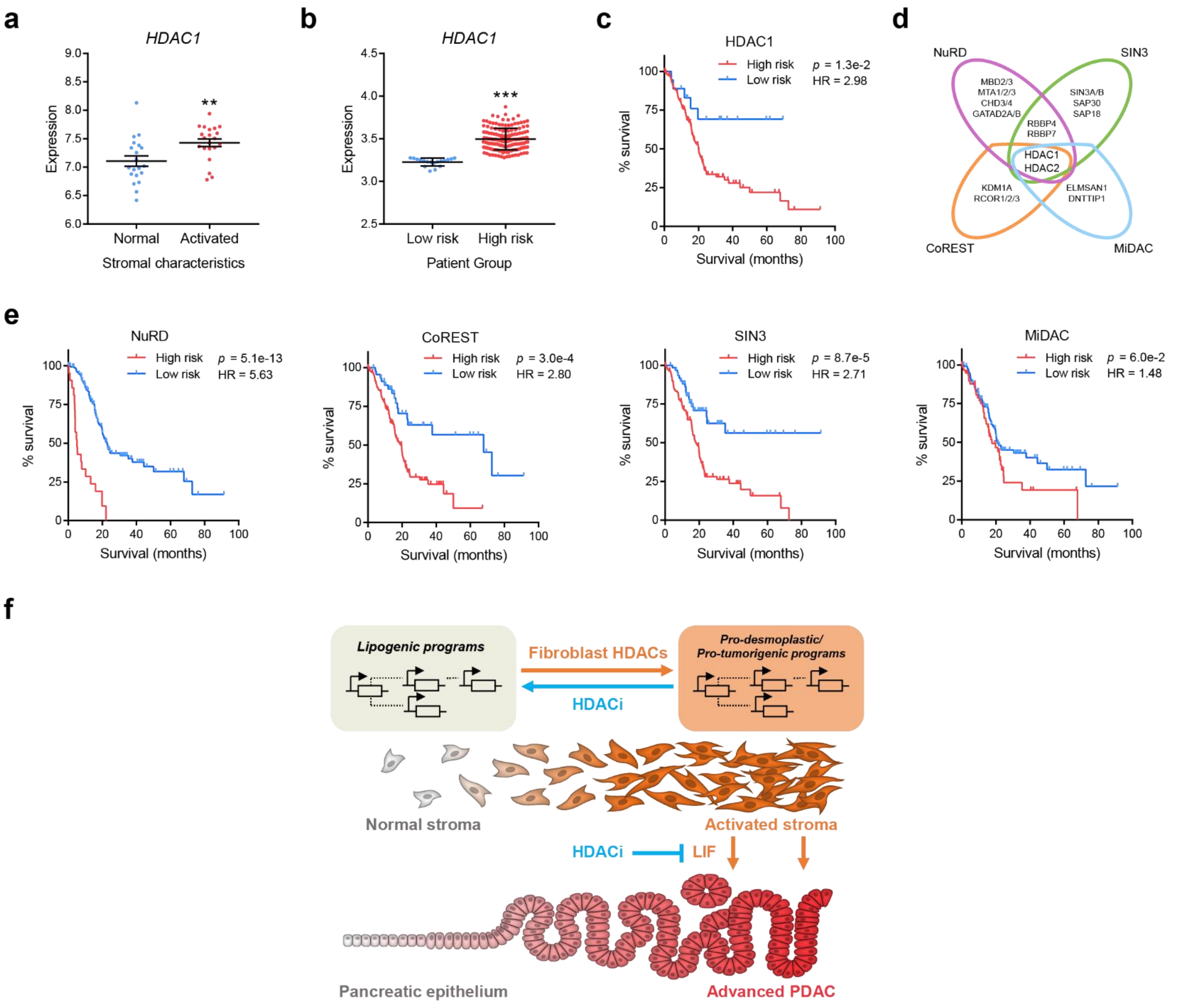


**Supplementary Fig. 10 | Correlations of patient prognosis with HDAC expression.**

**a** *HDAC1* expression in PDAC patient samples stratified by normal and activated stroma subtypes. **b, c** *HDAC1* expression in bulk tumor samples (**b**) and survival curves (**c**) from PDAC patients identified as high risk ($n$ = 158) or low risk ($n$ = 18) based on *HDAC1* expression. **d** Compositions of major HDAC1/2-containing complexes. **e** Survival curves of PDAC patients stratified by the expression patterns of HDAC1/2 complex components. NuRD, $n$ = 21 (high risk), 155 (low risk); SIN3, $n$ = 113 (high risk), 63 (high risk); CoREST, $n$ = 127 (high risk), 49 (high risk); MiDAC, $n$ = 60 (high risk), 116 (high risk). HR, hazard ratio. **f** Summary diagram highlighting the roles of fibroblast HDACs in promoting stromal activation and LIF-mediated pro-tumorigenic crosstalk by facilitating transcriptional program switch. Data in dot plots are presented as mean values ± SEM. ** $p < 0.01$; ***, $p < 0.001$. Two-sided t-test for *HDAC1* expression; log-rank test for survival analyses.

### Source Images

Uncropped images of the Western blotting data in Supplementary Fig. 6d. Arrowheads indicate target bands.

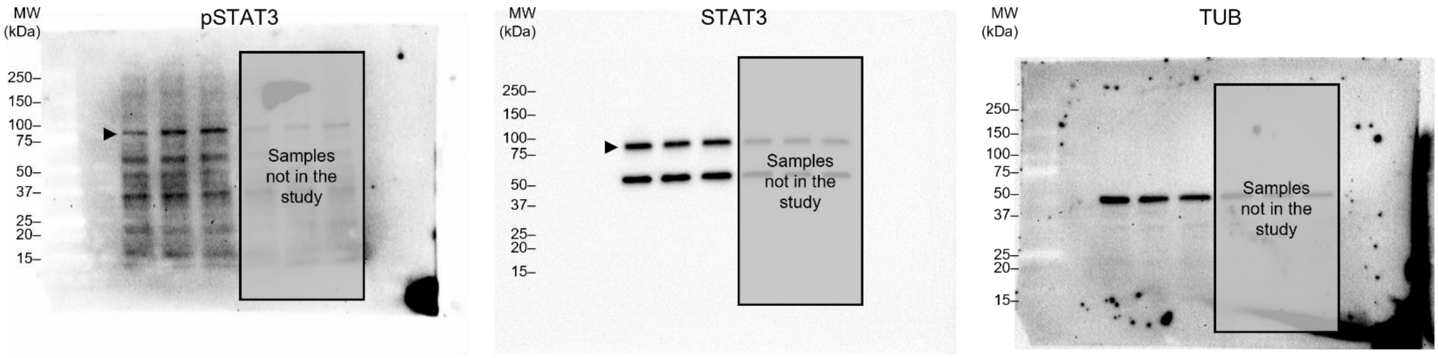
